# Biologically plausible mechanisms underlying motor response correction during reward-based decision-making

**DOI:** 10.1101/221770

**Authors:** Jung Hoon Lee

**Affiliations:** Allen Institute for Brain Science, Seattle, WA 98019

**Keywords:** reward-based decision-making, spiking network models, anterior cingulate cortex, prefrontal cortex, neural substrates of adaptive behaviors

## Abstract

Our propensity to acclimate to new surroundings and choose a goal-directed behavior for a maximal reward (i.e., optimal outcome) is natural, for it affects our survival. A line of studies suggested that anterior cingulate cortex (ACC) could be a potential hub for adaptive behaviors. For instance, an experimental study noted ACC’s contribution to selecting motor responses for maximal rewards; it found 1) that ACC neurons were selectively activated when the reward was reduced and 2) that suppression of ACC activity impaired monkeys’ ability to change motor responses to obtain the maximal reward. To probe ACC functions in adaptive behaviors, we sought biologically-plausible mechanisms to account for the experimental findings mentioned above by utilizing a computational model. Our simulation results raise the possibility that ACC can correct behavioral responses by reading out and updating the motor plans (guiding future motor responses) stored in prefrontal cortex (PFC).

## 1 Introduction

We are able to choose the best action to obtain the optimal outcome by evaluating available choices in terms of (expected) rewards; see [1, 2] for reviews. Our estimation of rewards’ values can change or evolve over time, and we must choose our actions strategically to gain maximal rewards. Then, how does the brain accomplish adaptive decision-making? In principle, the brain would need a precise mapping between at least three distinct factors, sensory inputs, (expected) rewards, and available choices. Notably, anterior cingulate cortex (ACC) has access to crucial information necessary for generating such mapping. First, dopamine, which has been thought to mediate reward signals [3, 4, 2], projects to ACC [5, 6]. Second, ACC interacts with PFC [7, 8] that is known to participate in executive functions and sensory signal processing [9, 10, 11, 12]. Indeed, ACC has been suggested to detect errors [13, 14, 15, 16, 17] and select a motor response for the maximal reward [18, 19, 19, 20]. These findings suggest that ACC can be a hub for adaptive behaviors or reward-based decision-making, but ACC’s exact functions remain poorly understood [21].

Here, we aim to better understand ACC’s contribution to adaptive decision-making by seeking biologically plausible mechanisms that can account for ACC activity during reward-based decision-making [20, 22]. A detailed account of ACC activity during reward-based decision-making was reported by Shima and Tanji [20], who trained monkeys to ‘push’ or ‘turn’ a lever when a visual cue was given. Monkeys received rewards when they selected the reward-choice, which was randomly chosen. After a block of trials, the reward-choice was changed from one to another; to monkeys, this transition was signaled by the reward reduction. When the reward was reduced, monkeys needed to switch from a previously selected motor response to another to obtain the maximal reward. During reward-based decision-making, the reward reduction depolarized ACC neurons, and suppressing ACC neuron activity impaired monkeys’ ability to switch motor responses accurately [20]. They further found that most ACC neurons responding to the reward reduction were sensitive to the sequence of a prior-response (before the reward reduction) and a post-response (after the reward reduction). Specifically, a subset of ACC neurons was activated when monkeys changed their responses from push to turn, but it remained quiescent when monkeys’ response sequence was reversed, indicating that ACC neurons can access information regarding identities of prior-responses and rewards and may use them to correct errors.

However, a more recent study [22] questioned the accuracy of ACC’s central role in error correction [20] by reporting that the lesion in ACC impaired monkeys’ ability to maintain the right choice rather than their ability to change their choice accordingly [22]. This inconsistency is puzzling because both studies used the same experimental protocol. To investigate the reasons behind the conflicting observations, we asked if a single network can account for both findings in our study. Specifically, we built a computational model consisting of ACC, prefrontal cortex (PFC), and motor cortex (MC) and mimicked the experimental protocol used to measure ACC activity [20]. To model the effects of dopamine in ACC, we simulated the effects of *D*_2_ receptors known to induce inhibition in ACC [23] by providing inhibitory currents to ACC neurons. In our model, when the reward is reduced, inhibitory currents are removed.

In the simulations, we found that 1) the motor response in MC was changed when the reward is reduced and 2) the removal of some ACC neurons can randomize motor responses. That is, our network model can explain both experimental observations [20, 22]. This model can be a starting point from which we can develop biologically-plausible models of ACC that can account for its diverse functions in a single framework [24, 7, 21, 25].

## 2 Methods

Our model consists of three cortical areas interacting with one another via inter-areal connections shown in Fig. 1 and Table 1. In each area of the model, neurons are split into several populations and have distinct connectivity. In addition, external background inputs (Poisson spike trains) are introduced to them to regulate their excitability and make their activity stochastic. The rates of Poisson spike trains are specific to neuron types (Table 2).

**Table 1:** For each pair of pre- and post-synaptic neurons, connections are drawn randomly using the connection probability and strength listed below. Each connection type was separated by semicolon in the table and has a conduction delay. The connections among neurons in the same area have 2.0 ms delay, whereas the connections across areas have 5.0 ms delay. To distinguish between MC and PFC, 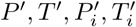 represent populations in MC, whereas *P, T, P*_*i*_, *T*_*i*_ represent them in PFC.

| Type | Param | Type | Param |
| --- | --- | --- | --- |
| $PT \rightarrow PT; TP \rightarrow TP$ | (0.1, 80) | $T \rightarrow P_i; P \rightarrow T_i$ | (0.1, 200) |
| $PT, TP \rightarrow NS$ | (0.1, 100) | $T_i \rightarrow T, T_i; P_i \rightarrow P, P_i$ | (0.2, -400) |
| $NS \rightarrow NS$ | (0.2, 60) | $T' \rightarrow T', T'_i; P' \rightarrow P', P'_i$ | (0.1, 50) |
| $NS \rightarrow PT_i, TP_i$ | (0.1, 100) | $T'_i \rightarrow T', T'_i; P'_i \rightarrow P', P'_i$ | (0.3, -400) |
| $PT_i \rightarrow PT; TP_i \rightarrow TP$ | (0.05, -100) | Connections across areas | |
| $PT_i \rightarrow PT_i; TP_i \rightarrow TP_i$ | (0.05, -50) | $TP \rightarrow T_i; PT \rightarrow P_i$ | (0.3, 100) |
| $T \rightarrow T; P \rightarrow P$ | (0.1, 150) | $P \rightarrow PT; T \rightarrow TP$ | (0.1, 3) |
| $T \rightarrow P; P \rightarrow T$ | (0.05, 50) | $T \rightarrow T'; P \rightarrow P'$ | (0.05, 10) |
| $T \rightarrow T_i; P \rightarrow P_i$ | (0.1, 20) | $T' \rightarrow TP, PT; P' \rightarrow TP, PT$ | (0.2, 100) |

**Table 2:** In the model, three different types of external inputs are used. First, background inputs are introduced to all neurons except NS neuron in ACC during the entire simulation periods. This table lists the rate of Poisson spike trains selected for neuron types. Second, visual cues are introduced to *P* and *T* populations (*P*′ and *T*′ in Table 1) in MC. Third, inhibitory currents are introduced to all ACC neurons to simulate the effect of *D*_2_ dopamine receptors [23]

| Type | Value |  |
| --- | --- | --- |
| Background (Hz) | ACC | $PT, TP = 500, PT_i, TP_i = 800$ |
| | PFC | $T, P, T_i, P_i = 800$ |
| | MC | $T, P = 300, T_i, P_i = 900$ |
| Visual Cue to MC (Hz) | $T, P = 300$ | |
| Inhibition via $D_2$ (pA) | $PT, TP, PT_i, TP_i NS = -250$ | |

**Figure 1:**
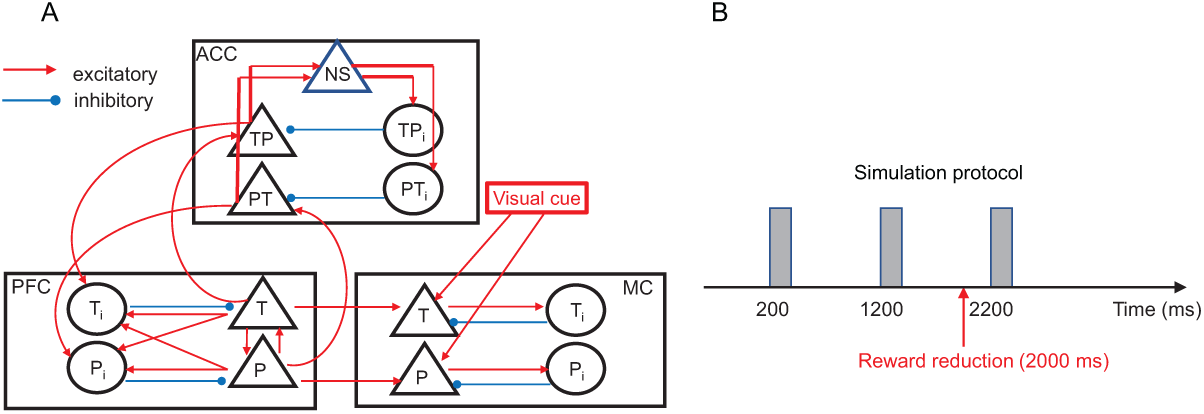
Structure of the model. (A), Schematics of the model consisting of three cortical areas. The triangles and circles represent excitatory and inhibitory neuron populations, respectively. The red and blue arrows indicate excitatory and inhibitory connections. For the sake of clarity, feedback connections from MC to ACC and recurrent connections within the same populations are not shown in the figure; see Table 1 for the full connectivity. (B), Protocol of simulations, three 200 ms-long visual cues are introduced at 200, 1200 and 2200 ms, and the reward is reduced at 2000 ms.

### 2.1 Neuron and Synapse models

All neurons in the model are implemented by leaky integrate-and-fire (LIF) neurons. Specifically, the subthreshold membrane potentials obey Eq. 1, and the crossing of the spike threshold (−55 mV) induces a spike. After the spikes are registered in the simulators, the membrane potentials are reset to the resting membrane potential (−70 mV), and no spike was allowed during the absolute refractory period (3 ms). The synapses in the model induce instantaneous voltage jumps in target neurons and decay exponentially according to Eq. 1.

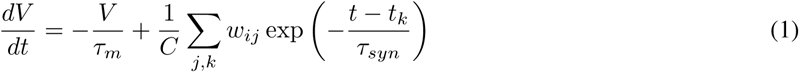

where *V* is the membrane potential; where *τ*_*m*_=10 ms and *τ*_*syn*_=2 ms are the time constants of membrane potentials and synaptic events, respectively; where *t*_*k*_ is the spike time of presynaptic neurons; where *w*_*ij*_ are synaptic strengths listed in Table 1; where *C*=250 pF is the capacitance of the membrane. The NEST’s native neuron model ‘iaf_psc_exp’ is used to implement this neuron [26]; see also [27] for additional information. The synaptic weights are fixed during the entire simulation; that is, we ignored the plasticity of synapses. The external background spike trains are mediated by synapses whose strengths are 200 pA.

### 2.2 Cortical area models

The model ACC consists of three excitatory and two inhibitory populations. The first two excitatory populations named *PT* and *TP* aim to simulate ACC neurons observed in the experiment, which were sensitive to the sequence of motor responses prior to and post the reward reduction [20]; *TP* and *PT* stand for ‘Turn to Push’ and ‘Push to Turn’, respectively. The two inhibitory populations *TP*_*i*_ and *PT*_*i*_ are exclusively connected to either *TP* or *PT* and provide feedforward inhibition onto the target population (Fig. 1). The third NS (non-selective) population receives excitatory inputs from both PT and TP and sends excitation to *TP*_*i*_ and *PT*_*i*_, which means that *NS* provides indirect feedback inhibition to *PT* and *TP*. ACC is known to have dopamine receptors, and we considered the effects of *D*_2_ receptors in ACC. As *D*_2_ receptors in ACC induce inhibition [23], we simulated the effects of dopamine by providing inhibitory currents to all ACC neurons (−250 pA). These inhibitory neurons are removed when the reward reduction is assumed.

The model PFC consists of two excitatory and two inhibitory populations. The two populations *P* and *T* are assumed to represent motor plans ‘Push’ and ‘Turn’, respectively. These neurons are connected with strong recurrent connections within *P* and *T* (Table 1) so that they could generate persistent activity which is believed to underlie working memory [28, 11, 29, 22]. *P* and *T* also interact with each other and with inhibitory populations (Fig. 1).

The model MC has a similar structure as that of PFC. As seen in Fig. 1, the two excitatory populations (*P* and *T*) interact with two inhibitory populations. *P* and *T* populations receive afferent inputs from *P* and *T* populations in PFC inspired by the influence of PFC on motor areas [12]. *P* and *T* are assumed to represent the actual motor response, and the activation of *P* (*T*) population is interpreted as Push (Turn) response. In addition, we assumed that these two excitatory populations receive afferent inputs elicited by visual cues; see [30] for the effects of sensory inputs onto motor cortex.

### 2.3 Simulation protocol

We mimicked the protocol used in the experiment (Fig. 1). Specifically, we simulated two trials (i.e., stimulus presentation) prior to the onset of the reward reduction and a single post trial. We performed simulations in two simulation conditions. In the reduced reward (RW) condition, the three visual cues are presented at 200, 1200 and 2200 msec, and the reward is assumed to be reduced at 2000 msec. Specifically, we removed inhibitory inputs to ACC neurons. We also performed the continuous reward (CW) as the control experiment, in which we maintained the constant reward level during the entire simulation period.

To examine the robustness of our results to stochasticity, 100 independent (but statistically equivalent) networks were instantiated using the same connectivity rule (Table 1), and 100 independent simulations were conducted. In each simulation, the background inputs to neurons in the model are independently generated using the rate given in Table 2.

### 2.4 Network implementation

We used NEST [26] to implement network models and simulate them. All neurons are identical in terms of internal dynamics, but they receive distinct external background and recurrent synaptic inputs Table1. Also, they receive external inputs representing background, visual cues and inhibition (mimicking dopamine *D*_2_ effects); the parameters are listed in Table(2).

### 2.5 Time course of PFC activity

To track changes in motor plans in PFC, we compared the outputs between *T* and *P* in PFC. Specifically, we split the spikes of PFC neurons into the 50 ms-long bins which do not overlap one another. Then, we calculated the selective index (*SI*) using the binned firing rates (*P*_*n*_, *T*_*n*_) of *P* and *T* populations using Eq. 2

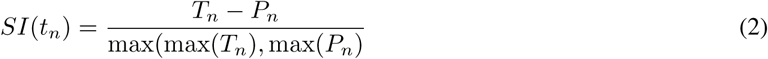

where *t*_*n*_ is the center of nth bin. When *SI* values are positive, *T* population produces stronger responses than *P* population. In contrast, *SI* values become negative, *P* population produces stronger responses. In the simulations, *P* and *T* populations are activated exclusively with one another due to the di-synaptic mutual inhibition (Table 1); once one of the populations is activated, its activity is persistent.

## 3 Results

Our model implements three cortical areas and their interactions to address our hypothesis. Each cortical area consists of identical LIF neurons and has multiple populations of excitatory and inhibitory neurons, which was inspired by cortical network models proposed earlier; see[31, 32] for instance. Here we assigned specific functions to the populations in each area. All these three areas interact with one another via connections illustrated in Fig. 1. Specifically, ACC and PFC are reciprocally connected [7], and MC projects nonspecific inputs to ACC. In all three areas, excitatory populations consist of 400 neurons, whereas inhibitory populations consist of 100 neurons; for clarity, the raster plots illustrated in Results show 10% of neurons. All connections are drawn randomly using the connection probabilities and synaptic weights listed in Table 1.

Inspired by earlier studies suggesting that PFC subserve working memory [28, 11, 29, 33] and that PFC and ACC are connected reciprocally [7, 8], we hypothesized 1) that PFC can hold motor plans in working memory to guide monkeys’ responses and 2) that ACC can read out and update these motor plans stored in PFC. With these hypotheses, we implemented PFC and ACC in the model. In addition, we implemented MC to simulate the full process of a motor response shift (i.e., lever movements) elicited by the reward reduction. Fig. 1A shows the schematics of our model. In ACC, three excitatory populations (*PT, TP* and *NS*) and two inhibitory populations (*PT*_*i*_ and *TP*_*i*_) interact with one another via connectivity listed in Table 1.

We simulated the experimental protocol with 3200 msec-long simulations by considering the effects of dopamine *D*_2_ receptors which are known to induce inhibition in ACC [23]; Fig. 1B illustrates the simulation protocol. In the simulation, the reward reduction is simulated by removing inhibitory currents to ACC (−250 pA). Each simulation consists of 3 blocks starting with a visual cue (Fig. 1B). As visual cues were used to signal monkeys to move the lever in the experiment [20], they must not carry any information regarding rewards. Therefore, in our model, we simulated visual cues with afferent inputs projecting to both populations in MC; see [30] for the effects of sensory inputs in motor cortex. As seen in Fig. 1B, visual cues are introduced to MC at 200, 1200 and 2200 msec after the onset of simulations, and their duration is 200 msec. To test reward effects on network activity, we performed simulations in two different conditions, continuous-reward (CW) and reduced-reward (RW). In the CW condition, the reward level remains constant, whereas in the RW condition, it is reduced at 2000 msec after the second visual cue. The initial motor plan is always ‘turn’ unless stated otherwise. As we assumed that PFC store motor plans using persistent activity, we provided 100 msec-long excitatory inputs to *T* population in PFC to initiate persistent activity within it before the first visual cue.

### 3.1 PFC can hold motor plan and guide MC responses accordingly

We first tested model responses in the CW condition (i.e., the high level of reward through simulations). As seen in Fig. 2A, *T* population in PFC, stimulated by external inputs before the first visual cue, fires continuously at a high firing rate due to strong recurrent connections in PFC (Table 1). That is, ‘turn’ is retained as the motor plan during the entire simulation period. With this motor plan in PFC, *T* population in MC responds to all three visual cues (Fig. 2B); ACC neurons remain quiescent. In the model, MC neurons are set to fire only when they receive afferent inputs from both PFC and visual cues. This makes MC neurons fire only during the visual cue, even though PFC neurons continuously project afferent inputs to MC *T* population. To examine if this result is robust over random connections and background inputs, we instantiated 100 independent networks. Each network has randomly chosen connections which are drawn from the same connectivity rule (Table 1), and we repeated the same simulation with independent sets of background Poisson spike trains (Table 2). In each simulation, we compared the average firing rates of *P* and *T* populations in MC during the presentation of third visual cue (2200-2400 ms). As seen in Fig. 2C, MC responses to the third are ‘turn’ in all 100 simulations, indicating that this network model sustains the same responses when the reward is maintained at a constant level.

**Figure 2:**
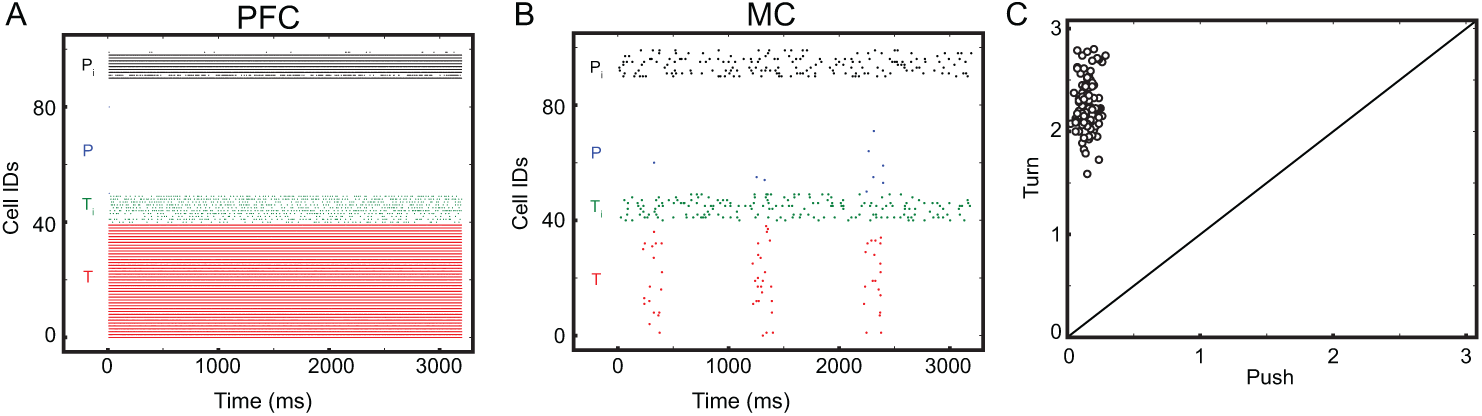
Model responses in the CW condition. For clarity, we show spikes from 10% of neurons. (A), Spikes from *T, P, T*_*i*_ and *P*_*i*_ populations of PFC are shown in red, blue, green and black, respectively. (B), Spikes from *T, P, T*_*i*_ and *P*_*i*_ populations of MC are shown in red, blue, green and black (C), Comparison between *T* and *P* populations of MC in 100 independent simulations. x- and y-axis represent the firing rates (Hz) of *P* and *T* populations during the presentation of third visual cue; *T* and *P* represent ‘turn’ and ‘push’. In each simulation, we calculated the average firing rate of *T* and *P* populations in MC. Each dot represents comparison between them.

### 3.2 ACC neurons update motor plans in PFC when they are depolarized by the reward reduction

We next tested responses of the model in the RW condition, in which the reward is reduced at 2000 ms (after the second visual cue). As in the CW condition, *T* population in MC responds to the first two visual cues (200-400 and 1200-1400 msec), but *P* population responds to the third visual cue (2200-2400 msec) (Fig. 3A). This means the motor response is updated successfully by the reward reduction. Then, how does the reward shift motor responses? In the model, when the reward is reduced, inhibition onto ACC neurons is lifted, making ACC neurons more responsive to afferent inputs. Among *TP* and *PT* populations in ACC, *TP* population is activated selectively (Fig. 3B) due to the excitation from *T* population in PFC (Fig. 3C).

**Figure 3:**
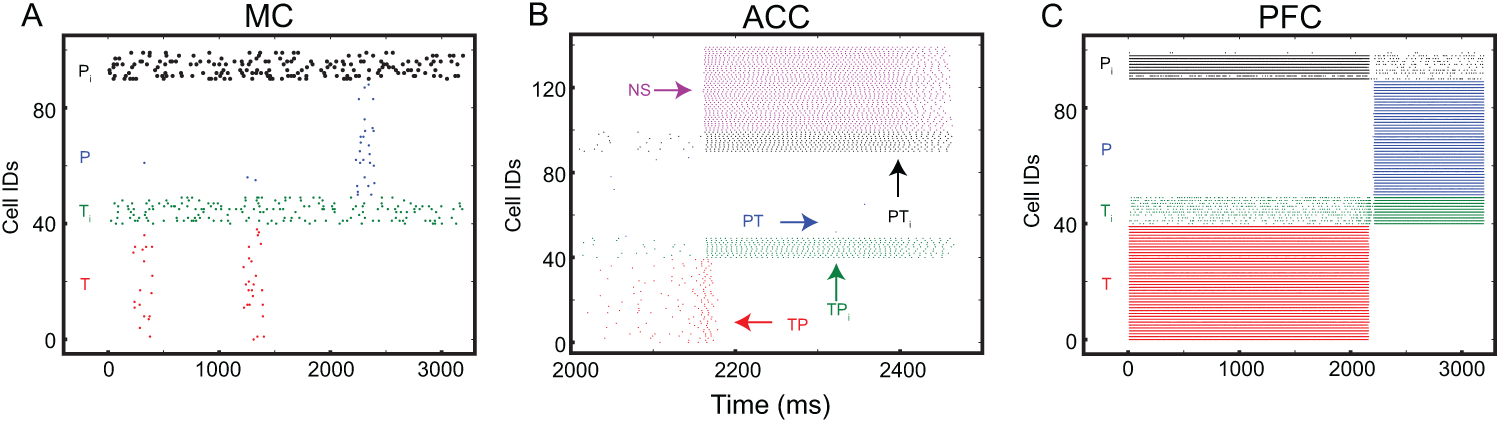
Model responses in the RW condition. For clarity, we show spikes from 10% of neurons. (A), Spikes from *T, P, T*_*i*_ and *P*_*i*_ populations of MC are shown in red, blue, green and black. (B), *T, P, T*_*i*_ and *P*_*i*_ and *NS* populations of ACC are shown in red, blue, green, black and magenta. (C), *T, P, T*_*i*_ and *P*_*i*_ of PFC are shown in red, blue, green, black.

We note that the activation of *TP* population in ACC initiates a series of changes in the model. First, the persistent activity of *T* population is turned off (Fig. 3C). Once *TP* population is active, it increases excitatory inputs to *T*_*i*_ in PFC, which in turn inhibits *T* population in PFC. Second, *P* population in PFC is turned on (Fig. 3C). As soon as *T* population in PFC is turned off, *P* population is liberated from lateral inhibition induced indirectly by *T* population (Fig. 1A) and starts firing due to external background inputs (Table 2); background inputs alone can drive PFC neurons to fire. Third, *NS* population in ACC becomes active due to excitation from *TP* population (Fig. 3B), which increases inhibition onto *PT* and *TP* by innervating *PT*_*i*_ and *TP*_*i*_. This delayed feedback inhibition suppresses *TP* activity after active firing, meaning the activation of *NS* population makes *TP* population transient, as observed by Shima and Tanji [20].

With these changes, *P* population in PFC is active, when the third visual cue is given (2200-2400 msec), and *P* population in MC becomes active consequently (Fig. 3A). After the switch in this response is instantiated properly, we assumed that the reward level is restored and re-introduced inhibitory inputs to ACC, making ACC neurons quiescent again (Fig. 3B). These results suggest that ACC can shift motor responses indirectly by updating the motor plan in PFC. As in the CW condition, we repeated simulations using 100 independently instantiated networks. To examine when the motor plan is switched in all these simulations, we calculated the selective index (*SI*) defined by Eq. 2 in each simulation. *SI* becomes positive when *T* population produces stronger responses in the selected 50 ms-time bin than *P* population. They are normalized to be between −1 and 1. The simulation results are summarized in Fig. 4. *TP* neurons in ACC generate stronger spiking activity than *PT* neurons do (Fig. 4A), the persistent activity in PFC is successfully transferred from *T* to *P* after the reward reduction (Fig. 4B), and motor responses are properly switched in all 100 simulations (Fig. 4C).

**Figure 4:**
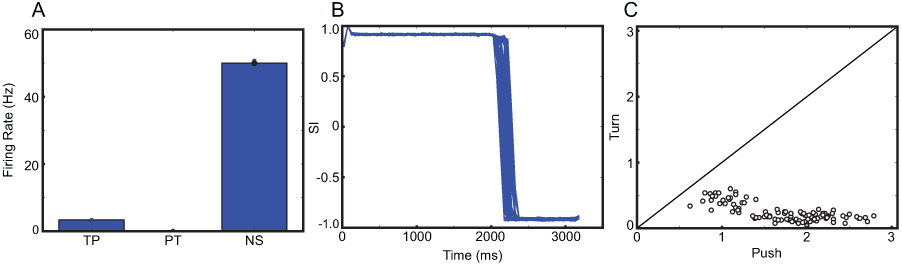
Responses from 100 independent simulations in the RW condition. (A), Mean values and standard errors of firing rates of *TP, PT* and *NS* populations in ACC neurons from 100 independent simulations; the spikes are collected from all ACC neurons during the entire simulation. In each simulation, we calculated the average firing rates of the three populations and the mean and standard errors over 100 simulations. (B), *SI* values of PFC from 100 populations. (C), Comparison between *T* and *P* populations in MC. x- and y-axis represent the firing rates during the presentation of the third visual cue. In 100 independent simulations, we calculated the average firing rates of T and P populations of MC. Each dot shows the firing rate in each simulation.

We further tested the importance of ACC in shifting motor responses in the model by removing inter-areal connections from ACC to PFC. As seen in Fig. 5A and B, *T* population in MC responds to the third visual cue when inter-areal connections from ACC to PFC are removed, confirming that it is ACC that corrects motor responses via interactions with PFC. Finally, we tested whether these results depend on the initial motor plan by changing it from ‘turn’ to ‘push’. As seen in Fig. 6, we observed the equivalent results even when the initial motor plan was switched.

**Figure 5:**
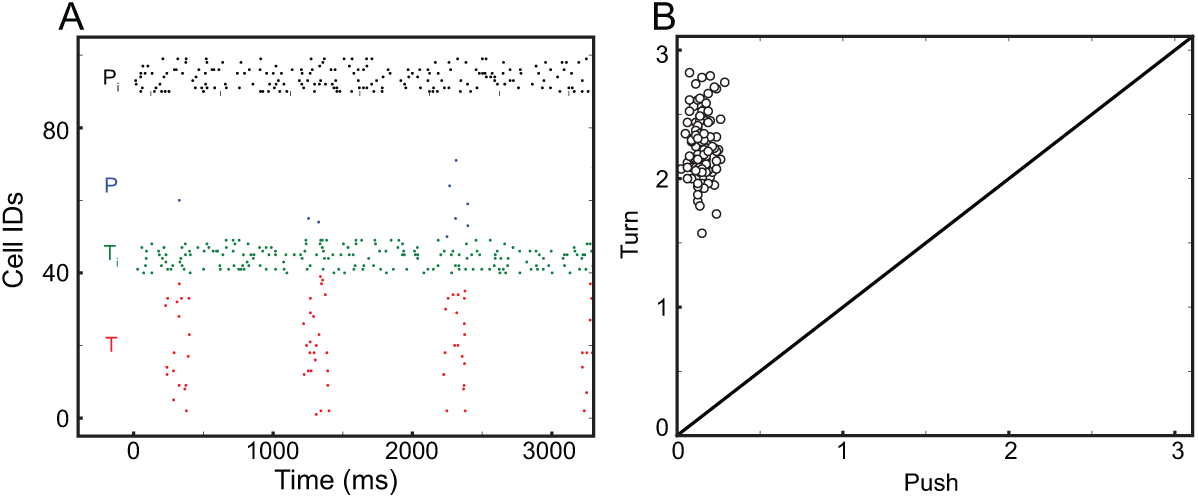
Model responses without afferent inputs from ACC to PFC. (A), Spikes from 10% of MC neurons. (B), Comparison between MC responses in the 100 independent simulations.

**Figure 6:**
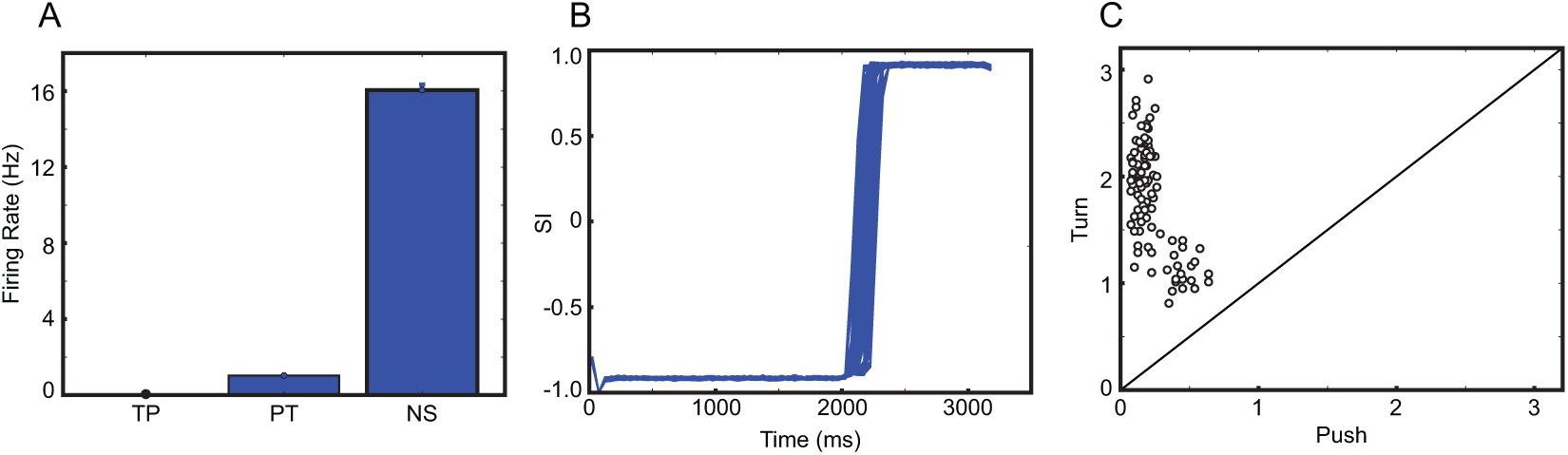
Model response with ‘Push’ as the initial motor plan. The same as in Figure 5, but a ‘push’ as the initial motor plan.

### 3.3 NS population is necessary for keeping motor plan in PFC stable

*NS* population in ACC shows delayed activity (Fig. 3B) and makes *TP* (*PT*) population activity transient. These transient and delayed responses are consistent with ACC activity patterns observed during reward-based decision-making [20], suggesting that *NS* population is necessary for replicating ACC activity patterns experimentally observed. To better understand *NS*’s function, we removed *NS* from ACC and repeated simulations in the RW condition. We found that MC responses to the third population became erroneous (Fig. 7A). In 6 out of 100 independent simulations, *T* population produces more responses than *P*; that is, motor responses are not properly switched in 6 simulations. Interestingly, we note that both *TP* and *PT* populations become active when *NS* population is removed (Fig. 7B). *PT* also becomes active because *P* provides excitation to *PT* population once the motor plan is shifted to ‘push’ (Fig. 7C). With *TP* and *PT* active, both *P* and *T* populations in PFC receive inhibition, disrupting the persistent activity in both populations (Fig 7C). After the quiescent period, the persistent activity can emerge either in *P* or *T* population (Fig. 7D).

**Figure 7:**
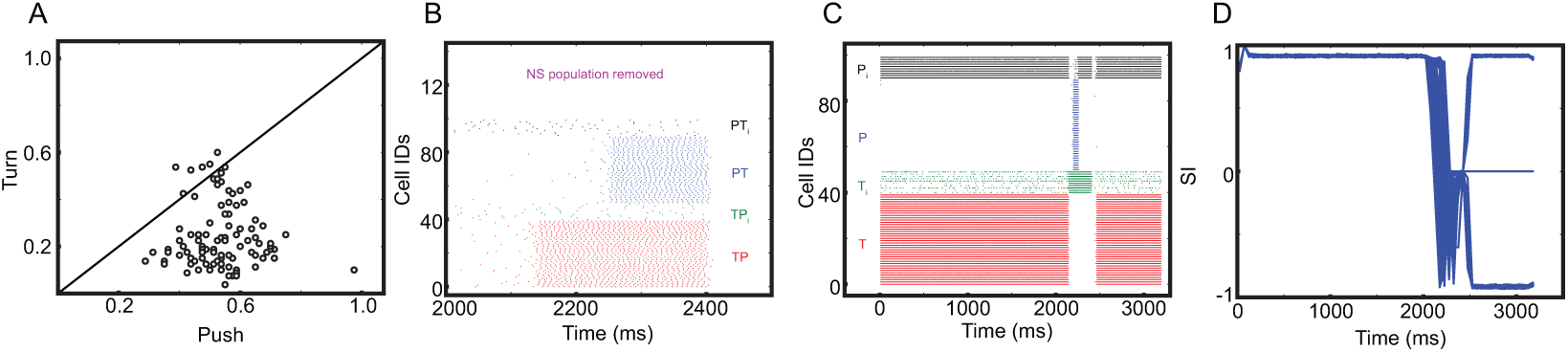
Model responses without *NS* population. (A), Comparison between MC responses in the 100 independent simulations. (B), Spikes from 10% of ACC neurons. *TP, PT, TP*_*i*_, *PT*_*i*_ and *NS* populations of ACC are shown in red, blue, green, black and magenta. (C), Spikes from 10% of PFC neurons. *T, P, T*_*i*_, and *P*_*i*_ of PFC are shown in red, blue, green, black. (D), Time course of *SI*.

These observations led us to hypothesize that *NS* population may be responsible for stabilizing the motor plan stored in PFC. To test this hypothesis, we prolonged each simulation to have 6 visual cues. Specifically, 3 more visual cues are added, and simulations before the fourth visual cue are indeed identical to those with 3 visual cues discussed above. Figure 8A shows how PFC activity evolves over time in the RW condition without NS population. As seen in the figure, the persistent activity moves back and forth after the reward reduction. This unstable PFC activity makes the MC responses to the last four visual cues erroneous; *T* population, instead of the correct *P* population, sometimes responds to visual cues (Figs. 8B-E). We also note that incorrect answers to the first cue (after the reward reduction) are less frequent than those to other three cues (Fig. 8B-E). These results are consistent the observation that the lesion in ACC disrupted monkeys’ ability to maintain the correct choice [22], while it did not strongly degrade the accuracy of responses to the first cue (after the reward reduction).

**Figure 8:**
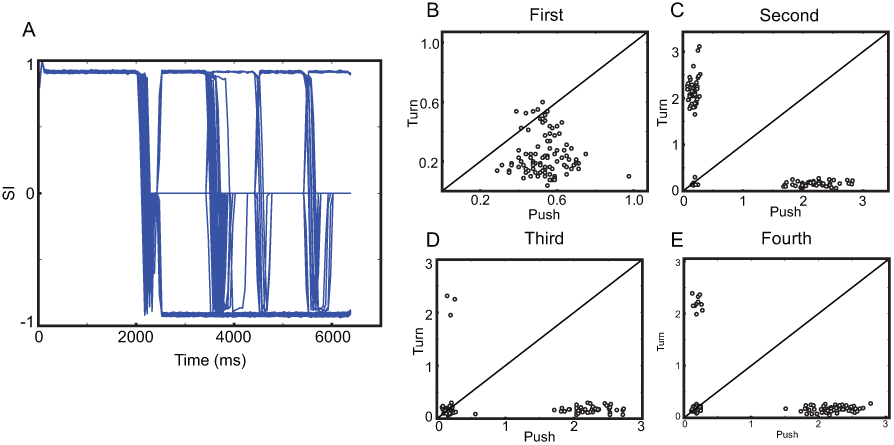
Model responses to 6 visual cues without NS population. (A), Time course of *SI*. (B)-(E), Comparisons between MC responses to the last four visual cues. The panels ‘First’, ‘Second’, Third’, and ‘Fourth’ above indicate the third, fourth, fifth and sixth visual cues in the simulation, respectively.

## 4 Discussion

To gain insights into ACC’s function in regulating motor responses during adaptive behaviors (and reward-based decision-makings), we sought neural circuits capable of 1) replicating ACC neuron activity patterns observed during a simple reward-based decision-making [20] and 2) explaining the effects of lesions in ACC in monkeys’ behaviors [22]. In our model, there are two types of excitatory ACC neurons, *PT/TP* and *NS*. They show transient and delayed responses, which is experimentally observed [20]. Our simulation results suggest that the two neuron types have distinct functions. *TP/PT* neurons can read out and update motor plans when the reward is reduced, which is consistent with correlated activity in the two areas [34, 7] and the suggested central role of PFC in error detection [35]. In contrast, *NS* neurons, internally driven by *PT/TP* neurons, do not participate in correcting responses directly. Instead, they enable MC and PFC to maintain the right choice by suppressing *TP/PT* neuron activity. Thus, our model raises the possibility that ACC can update and stabilize motor responses, which, in turn, can explain the two seemingly conflicting experimental findings [20, 22] with a single mechanism.

Below, we discuss the potential links to the earlier models of ACC; see [21] for a comprehensive review.

### 4.1 ACC as a conflicting monitor

In our model, populations in ACC have exclusive counter parts in PFC; for instance, *PT* (*TP*) population in ACC does not interact with *T* (*P*) population in PFC (Fig. 1A). These exclusive connections can be ideal to obtain maximal rewards during reward-based decision-making [20]. In the brain, however, the exclusive connections may be too restrictive, and the weak cross connections from PFC to ACC (e.g., *P* in PFC projects to *TP* in ACC) exist likely. With these cross connections, our model can effectively account for conflicting monitoring attributed to ACC [36, 17], if PFC can encode multiple aspects of sensory evidence (e.g., color and identity of word). With PFC neurons responding to both the colors and identities of words, a word ‘red’ written in blue color will innervate more PFC neurons than a word ‘red’ in written in red color. That is, the incongruent word will enhance ACC activity more than the congruent one, as the conflicting monitoring theories suggested [36, 17], which is supported by PFC’s dominant role in action monitoring occurring in ACC [35].

### 4.2 Extension to unified framework of ACC’s function

It is well understood that ACC activity is elicited during various types of cognitive tasks [24, 7, 21, 25]. For instance, ACC activity is reported to be correlated with expected values of outcomes [37, 22], and ACC can also encode the efforts for given tasks or choice difficulty [38, 39, 40, 41, 42]. Although the diverse hypothetical functions of ACC pose great challenge when developing a unified theory of ACC’s functions, recent theoretical studies provide potential mechanisms to account for their diverse functions in a single framework; see [21] for a review. The predicted outcome model (PRO) and reward value prediction model (RVPM) assume that ACC neurons predict the value or the likelihood of future outcomes via reinforcement learning [43, 44, 21].

Based on these studies, it seems natural to assume that behavioral tasks with stochastic rewards (i.e., uncertainty of rewards) are necessary to fully understand ACC function. As our goal was to elucidate biologically-plausible mechanisms to account for a simple deterministic behavioral task, in which components of the task have no uncertainty, we did not consider ACC functions associated with stochastic rewards in this study. In the future, we will extend our model by considering 1) more complex tasks with stochastic rewards (i.e., uncertainty) and 2) *D*_1_ receptor which was suggested to underlie ACC’s contribution to effort-based decision-making [41]. The extended study could bring us a step further to a single biologically-plausible theory that can account for diverse functions of ACC. It should be noted that biologically-plausible models can simulate multimodal neural signals including functional magnetic resonance image, electroencephalogram and single unit activity. As these multimodal signals can be used to test a theory against abundant sets of data, we expect the extended version of our biologically-plausible model to provide rigorous ways to test multiple theories regarding ACC’s functions. For instance, it will produce multiple scale neural responses derived from PRO and RVPM models, and such predictions will help us choose the model that describes ACC’s functions more accurately.

## Acknowledgements

We thank the Allen Institute founder, Paul G. Allen, for his vision, encouragement and support.

